# Utilization of high throughput genome sequencing technology for large scale single nucleotide polymorphism discovery in red deer and Canadian elk

**DOI:** 10.1101/027318

**Authors:** Rudiger Brauning, Paul J Fisher, Alan F McCulloch, Russell J Smithies, James F Ward, Matthew J Bixley, Cindy T Lawley, Suzanne J Rowe, John C McEwan

## Abstract

Deer farming is a significant international industry. For genetic improvement, using genomic tools, an ordered array of DNA variants and associated flanking sequence across the genome is required. This work reports a comparative assembly of the deer genome and subsequent DNA variant identification. Next generation sequencing combined with an existing bovine reference genome enabled the deer genome to be assembled sufficiently for large-scale SNP discovery. In total, 28 Gbp of sequence data were generated from seven *Cervus elaphus* (European red deer and Canadian elk) individuals. After aligning sequence to the bovine reference genome build UMD 3.0 and binning reads into one Mbp groups; reads were assembled and analyzed for SNPs. Greater than 99% of the non-repetitive fraction of the bovine genome was covered by deer chromosomal scaffolds. We identified 1.8 million SNPs meeting Illumina *InfiniumII* SNP chip technical threshold. Markers on the published Red x Pere David deer linkage map were aligned to both UMD3.0 and the new deer chromosomal scaffolds. This enabled deer linkage groups to be assigned to deer chromosomal scaffolds, although the mapping locations remain based on bovine order. Genotyping of 270 SNPs on a Sequenom MS system showed that 88% of SNPs identified could be amplified. Also, inheritance patterns showed no evidence of departure from Hardy-Weinberg equilibrium. A comparative assembly of the deer genome, alignment with existing deer genetic linkage groups and SNP discovery has been successfully completed and validated facilitating application of genomic technologies for subsequent deer genetic improvement.

## INTRODUCTION

New Zealand has the largest farmed deer industry in the world; trophy hunting and velvet products exist, however venison is the primary export product with 1.2 million carcasses being exported, primarily to Europe, in 2009 (Deer Industry New Zealand). New Zealand (NZ) has no indigenous ruminants and there have been many importations of deer into NZ over the last 120 years. Most NZ deer are considered to fall into three “breeds”, Western European, Eastern European and Canadian Elk. Here, the term “breed” is used to define the three groups, although they include animals that have been classified as different *Cervus elaphus* sub-species. The Western European group is derived from Scotland as well as from English reserve parks (which in themselves sourced red deer from external sources such as Germany); they are likely to primarily include *C. e. atlanticus* and/or *C. e. scoticus.* The Eastern European stock was imported primarily from Hungary, the former Yugoslavia and Romania and is assumed to comprise *C. e. hippelaphus* genetics. The elk (*C. e. canadensis*) was originally sourced from Elk Island in Alberta, Canada.

Livestock industries including those for cattle and sheep utilize SNP markers for a variety of purposes, such as parentage assignment and for ascertaining levels of admixture or breed composition (Fisher et al. 2009; Kuehn et al. 2011; Frkonja et al. 2012). SNP chips have been especially useful in generating relationship matrices to enable prediction of genetic performance via genomic selection (GS) (Meuwissen et al. 2001; Hayes and Goddard 2008). In the dairy industry, where uptake of SNP chip technology is high, new breeding systems are being designed to incorporate genome-assisted selection (De Roos et al. 2011; Pryce and Daetwyler 2012). Several catalogues now indicate whether genomic information has contributed to the ranking of bulls; for example, see Holstein Canada’s Lifetime Profit Index, LPI rankings (Holstein Canada Home Page). Because GS offers improved genetic gain for both well-measured and sex-dependent, late- or hard-to-measure traits, it could provide great value to the New Zealand deer industry. However, GS cannot be achieved without first putting together a large-scale marker discovery program. The sheep genomic platform initially used the bovine genome to aid its sequence assembly and SNP discovery outcomes and has used comparative genomics further, also enlisting the dog and human genomes in order to build a virtual sheep genome (Dalrymple et al. 2007). An ovine SNP discovery program resulted in the building of an ovine 50K SNP chip which has subsequently been used for genomic applications including the description of the genetic structure of sheep breeds (Kijas et al. 2012). This report describes the generation of 28 billion base pairs (Gbp) of cervid sequence and the subsequent use of the bovine genome to aid the sequence assembly process, ultimately for SNP discovery in deer.

## MATERIALS AND METHODS

### Animal Selection

Seven deer were selected for sequencing, balancing a variety of factors including sampling the genetic diversity of the species in NZ, sex, DNA sample availability, access to pedigrees and New Zealand industry impact.

The animals selected are shown in Table 1; the *Cervus elaphus* species’ natural habitat covers a wide geographic range across the United Kingdom and Europe, Asia and North America. The seven selected animals represent sub-species from the United Kingdom, Europe and Canada and their pedigrees trace back to importations from the geographical locations described in Table 1. It should be noted that the English Park deer, whilst referred to as *“C. e. scoticus”,* may have historical English, Scottish and/or German genetic influence. Six of the seven animals were female, ensuring good coverage of the X chromosome. The seventh animal (EAS1) was an influential industry stag and provided some low level sequence coverage of the Y chromosome.

**Table 1.** The geographic sources of the animals selected for sequencing. The seven deer that were sequenced (individuals) were sourced within New Zealand. They included six females (F) and one male (M). The animals’ recorded years of birth (Y OB) are given and their pedigrees traced their ancestors back to the exotic locations described. The seven deer belong to three sub-species, *C. e. canadensis* (elk), *C. e. hippelaphus* (Eastern European red deer) and *C. e. scoticus* (Western European red deer).

| *Individuals* | *Sex* | *YOB* | *Geographic ancestral origins* | *Sub-species* |
| --- | --- | --- | --- | --- |
| ELK1 | F | 1990 | Elk Island National Park, Alberta | *C. e. canadensis* |
| ELK2 | F | 1985 | Elk Island National Park, Alberta | *C. e. canadensis* |
| HUN1 | F | 1994 | ¾ Hungarian, ¼ Yugoslavian | *C. e. hippelaphus* |
| WOB1 | F | 2008 | English Park (Woburn Abbey, UK) | *C. e. scoticus* |
| WAR1 | F | 2008 | English Park (Warnham Estate, UK) | *C. e. scoticus* |
| RED1 | F | 1997 | Invermark, Scotland | *C. e. scoticus* |
| EAS1 | M | 1997 | ¼ Hungarian, ¾ Yugoslavian | *C. e. hippelaphus* |

Deer farming began approximately 30 years ago in New Zealand both from live capture of wild animals and importation. The pedigrees, provided in Additional file 1, consist of less than six generations of controlled breeding, assisted by parentage verification using microsatellite markers. With the exception of RED1, the animals’ ancestors can be traced back to the country they were imported from. Five of the pedigrees show no documented evidence of inbreeding, however there is co-ancestry in HUN1 and also in the industry stag EAS1. Co-ancestry was determined for HUN1 and EAS1 using PROC INBREED, SAS. Despite having the “YUG1” stag in its pedigree multiple times, EAS1 was estimated to be just 3.1% inbred. HUN1 and EAS1 also share some common ancestry (15.2%), which in terms of SNP discovery represents a loss of ~1 of the 14 genomes sampled in the project.

DNA was extracted from semen according to company guidelines for PhaseLock Gel™ tubes (Qiagen Inc, CA, USA) or from blood (Montgomery and Sise 1990). DNA was deemed to be of good quality based on spectrophotometric measurements. Additionally, the samples were electrophoresed on 1% agarose to determine whether the DNA was of sufficient length for sequencing i.e. consisted of a single band of ~20Kbp or greater length. Finally, to confirm that the samples were from the expected animals, they were genotyped with the standard microsatellite deer parentage identification panel by a commercial laboratory (AgResearch, GenomNZ, NZ) prior to sending to Illumina Inc. (CA, USA) for sequencing._Spectrophotometric measurements confirmed that all of the DNA samples had A260/280 ratios > 1.8 i.e. they were of sufficient purity for sequencing. Further, the agarose gel confirmed that the DNA of all seven samples primarily consisted of fragments that are 20 kb or greater in length and that are consequently suitable for sequencing. And finally, parentage verification confirmed that all of the samples were from the expected animals.

### Sequencing and Assembly

A whole genome sequencing approach was undertaken using the GAII-X sequencing platform (Illumina Inc.). The methodology was based on a paired end approach with 2 x 100 bp reads (Illumina Paired-End Sequencing Information). In collaboration with Illumina, this was carried out according to standard procedures by the Illumina FastTrack Sequencing Services, Hayward, CA, USA. One lane of sequencing was carried out for each animal except “RED1” which was sequenced on two lanes. In-house scripts (Python) were used to analyze read length, quality values per base and sequence bias.

RepeatMasker (Smit et al. 1996-2010) was used with a custom repeat library to remove repetitive DNA from downstream analysis. This library consisted of *ruminantia* repeats from RepBase (Genetic Information Research Institute) supplemented with the reads that were present in more than ten copies per lane. Further, this was supplemented by read fragments that had thousands of hits against the cattle genome.

Mega BLAST 2.2.26 (Zhang et al. 2000) was used to map the deer reads to the cattle reference genome UMD 3.0 with settings as follows:

-D 3 –t 21 –w 11 –q -3 –r 2 –G5 –E 2 –s 56 –N 2 –F “m D” –UT

If a read had more than one hit, it was considered for mapping only if the second most likely hit had an adjusted e-value (divide e-value of best hit by e-value of second best hit) of 1e^−20^ or smaller (i.e. the 2^nd^ hit is considerably less likely than the first). After mapping, the bovine genome was divided into adjacent one Mbp bins and the mapped deer reads were placed into the corresponding bins. This enabled a “divide-and-conquer” approach to make the subsequent assembly less computationally demanding and use our compute farm efficiently. Where only one of the paired end sequences mapped to a bin, the partner read was added to the group.

Assembly parameters were first determined through simulation. A one Mbp region of the bovine genome was selected (Zimin et al. 2009) and Illumina 2 x 100 bp paired end reads with an insert size of 500 bp were simulated using the MetaSim package (Richter et al. 2008). Next, Velvet (Zerbino and Birney 2008) was used with different kmer sizes i.e. varying lengths of overlapping, shared nucleotide sequence comparisons. The optimal kmer size of 31 was determined by a combination of N50 contig length and percent coverage of the sample region by contigs. We detected a wide variation in insert size across our eight sequencing libraries. Because of the low coverage depth per library we couldn’t run the assembly on a per library basis with optimized settings but opted for an assembly of all libraries together and dropped paired-end information because it was not consistent across our libraries. Each one Mbp bin was assembled individually using Velvet 0.7.55 with the following settings: velveth 31 –short ; velvetg –cov_cutoff auto. After the assembly had taken place, the contigs and singletons were mapped back to their respective one Mbp bins on the cattle genome, using Mega BLAST with the following settings:

-m 4 –D 2 –t 21 –W 11 –q -3 –r 2 –G 5 –E 2 –s 56 –N 2 –F “mD” –U T –v 100000 – b 100000 –a 4.

A customized compiled BASIC program MELD (McEwan and Brauning: available upon request) was used to build scaffolds and transfer the order, orientation and spacing of contigs and singletons from the blast results. These were then collapsed into a single chromosomal scaffold consisting entirely of deer sequence but based on bovine chromosomal order. Once the deer chromosomal scaffolds were built they were masked again using RepeatMasker and the custom library described above.

To assess assembly quality we checked to what extent cattle nucleotide refseqs (retrieved from NCBI, January 2010) were covered by our assembly. We mapped masked cattle refseqs onto the deer assembly using Mega BLAST with the following settings: -D 3 –t 21 –W 11 –q -3 –r 2 –G 5 –E 2 –s 56 –N 2 –F “m D” – U T.

### SNP Detection

Once the deer chromosomal scaffolds were assembled, all of the deer reads were mapped to them using Mega BLAST with parameter settings:

-m 3 –D 2 –F “m D” –U T –v 100000 –b 100000 –a 2

Clonal reads (artefacts created during sequencing) were detected where reads start at the same base when mapped to the assembly. Where these occurred they were collapsed down to one copy. We analyzed the sequence depth across all of the repeat-masked chromosomal scaffolds and based on the shape of the read depth distribution discarded SNPs that were covered by too few or too many reads.

### Filtering SNPs for suitability as *Infinium II* SNP chip markers

The goal was to select SNPs that could become DNA markers and more specifically, could be incorporated into an *Infinium II* cervine SNP chip (Illumina). Only SNPs that had already passed the depth filter described above were considered. They were also required to contain at least 80 bp of non-repeat-masked sequence flanking both sides of the SNP. In the first of several filters, the SNPs that had one or more additional SNPs within 10 bp were removed; this was referred to as the *proximity filter.* Next, as the SNPs could either be genuine, or sequencing artefacts, they were placed into three classes: A) SNP heterozygous in multiple animals, B) multiple animals showing a total of two alleles (SNPs can be homozygous or heterozygous per animal), C) SNP heterozygous in a single animal all other animals do not differ from the reference. This is referred to as the *class* filter. Then, the A/T and G/C SNPs were removed because the *Infinium II* SNP chip technology does not utilize these polymorphisms; this was called the *nucleotide exchange* filter. And finally, the SNPs that remained after the filtering steps had been applied, and their flanking DNA were placed into the *Infinium II* design pipeline (Illumina).

### SNP marker Validation

A sample of 270 class A SNPs was converted into markers for use on a mass spectrophotometry platform (Sequenom). The SNPs were placed in eight multiplexes. The markers were selected, based on UMD3.0 position, for subsequent utility in population genetic and candidate gene analyses (unpublished). They were first characterized to confirm that the SNP filtering pipeline above had identified valid markers with utility for genomic applications. The source of the sequence (elk vs red deer) was not considered during SNP selection. Initially, the seven sequenced deer in addition to 90 other deer from a “breed reference” dataset were genotyped. The NZ-derived breed reference animals were previously classified as Western European (37 red deer, expected to have Scottish, English and German ancestry), Eastern European (21 red deer, expected to have Yugoslavian, Hungarian or Romanian ancestry), Canadian elk (25 elk, expected to have Canadian ancestry) or unknown breed (seven animals). The breed classification had been based on pedigree and microsatellite data (unpublished). Genotypes were produced according to standard Sequenom protocols and no optimization of the multiplexes was attempted. SNPs whose genotypes passed technical acceptance criteria (Sequenom “conservative” and “moderate” classes) for >90% of the tested samples were investigated further for polymorphism, error and for adhering to Hardy-Weinberg.

To complete the SNP quality control, six known sire-offspring pairings were genotyped. These pairings were used to test for mismatches, the presence of which would implicate genotyping error and/or false sequence-based SNP prediction.

### Assigning deer linkage groups to the bovine reference genome

Deer and bovine genetic maps exist (Slate et al. 2002; Ihara et al. 2004) and shared markers were previously used to show which chromosomes were syntenic (Slate et al. 2002). The procedures described below were carried out to assign the deer linkage groups to bovine reference genome positions and to confirm that the deer chromosomal scaffolds mapping procedures above were consistent with these alignments.

Next the deer genetic map was aligned to bovine sequence order. First, the primer sequences of microsatellites located on the deer genetic map were aligned to UMD3.0 and to the deer chromosomal scaffolds using BLASTN with the following settings: - F F -W 7 -m 9. Only markers whose primers both mapped within 400 bp of one another, and in opposite directions were considered for additional mapping analysis.

Restriction Fragment Length Variants, RFLVs, also located on the red deer genetic map (Slate et al. 2002) were mapped using MegaBLAST with the following parameters:

-F "m D" -U T -t 21 -W 11 -q -3 -r 2 -G 5 -E 2 -s 56 -N 2 -m 8

Finally, reciprocal MegaBLAST alignments were carried out; UMD3.0 sequences corresponding to the microsatellite and RFLV markers were mapped to the deer chromosomal scaffolds using the following parameters:

-F “m D” -U T -t 21 -W 11 -q -3 -r 2 -G 5 -E 2 -s 56 -N 2 -m 8

Chromosomal scaffold hits that mapped to expected locations according to published syntenic relationships were assigned to the appropriate deer linkage groups.

Assigning chromosomal scaffolds were also assigned to locations for linkage groups that had undergone “fission”. Six Pecoran ancestral chromosomes each split into two during the evolution of modern red deer. These fissions did not occur in the bovine genome; therefore six bovine chromosomes are syntenic to six pairs of deer chromosomes (Slate et al. 2002). For each bovine chromosome, the genetic markers that had mapped to the two syntenic deer linkage groups were aligned via Mega BLAST, using the conditions described above. Deer chromosomal scaffolds positions could not be assigned to linkage groups if they mapped to UMD3.0 in the region between the last marker of the proximally mapped linkage group and the first marker of the distally mapped linkage group i.e. the region within which the breakpoint is thought to exist. All other deer chromosomal scaffolds were assigned to the appropriate linkage group.

## RESULTS AND DISCUSSION

### Sequence outputs

A total of 284 million 100 bp reads (i.e. 28.4 Gbp) were generated, equating to 9x coverage in total for the seven-deer set. Figure 1 shows Phred scores across the entire length of the 100 bp reads. The mean Phred scores dropped from 33 at the start of the read to 16 at base 100. The Phred score above 15 was deemed to be adequate given that a depth of nine was achieved; therefore the reads were not generically trimmed and the entire 100 bp fragments were included in subsequent filtering stages.

**Figure 1.**
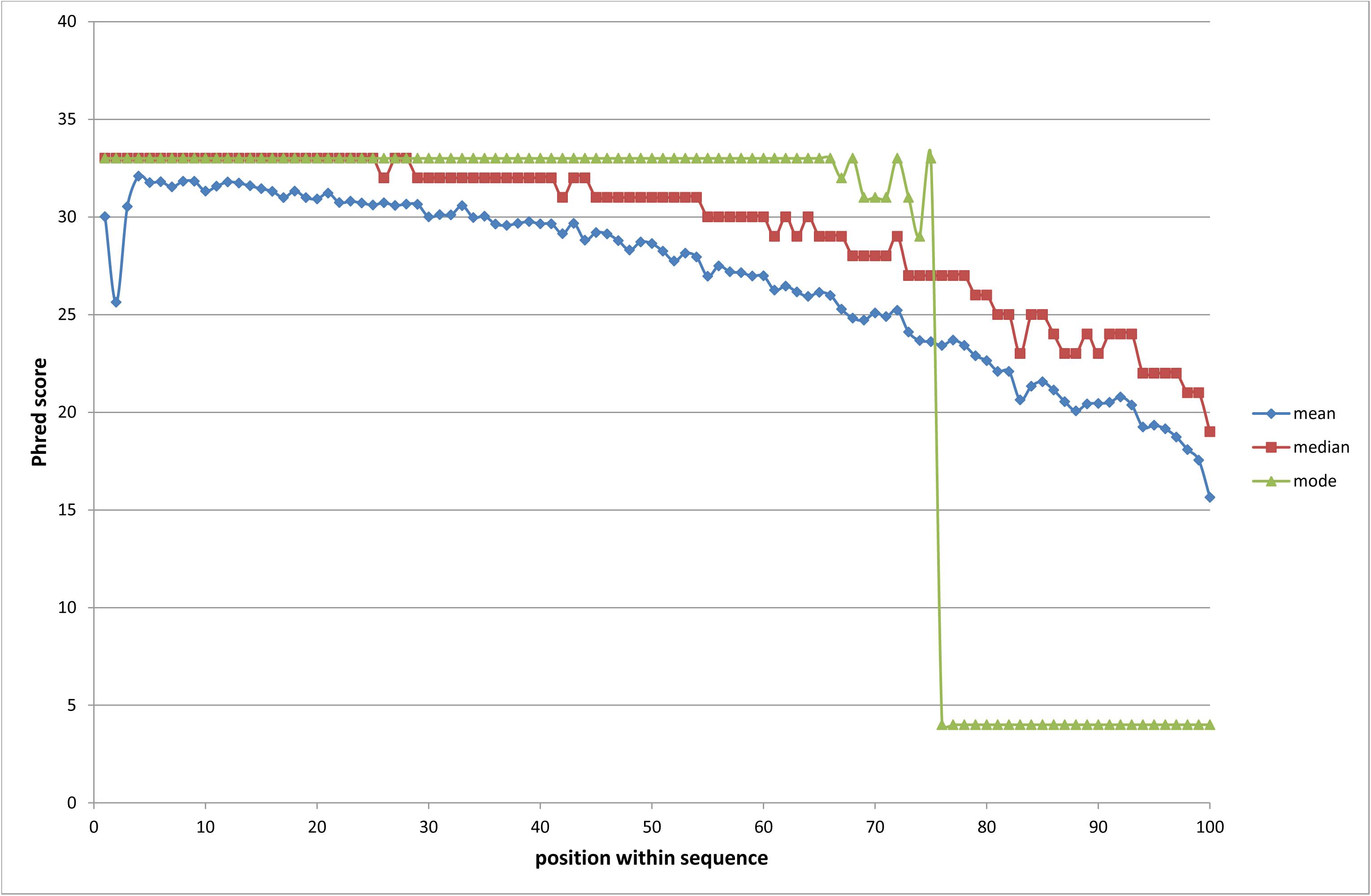
Sequence read quality (Phred) scores for each position across the 100 bp reads. The mean (blue), median (red) and mode (green) Phred scores for each of the 100 bases of the sequence reads. All three values dropped as the reads extended, such that the median Phred scores dropped to 18 for the last base.

### Masking Repetitive DNA

*The Ruminantia* repeat library alone allowed repeat-masking of 34% of the reads and these masked reads were masked to a level of 88% of their length on average. The custom library increased the percent of masked reads to 41% of all reads. These masked reads were masked to a level of 91% of their length. Sensitive masking of repeats was carried out to improve mapping specificity.

### Mapping to the bovine reference genome

In total 48% of the repeat-masked reads mapped to UMD3.0 (one or more hits) and 90% of all hits were unique. The mapping specificity was very high as only 0.1-0.4% of all read pairs mapped to multiple cattle chromosomes. In total, 99.97% of all mapped read pairs had the two reads mapped in the correct orientation.

Using mapping position information we deemed our insert sizes to average 200 bp, which is less than half of what was originally expected. The distributions of the distance between the inner ends of the paired end reads is shown in Figure 2 which shows approximately half of the pairs had some overlap. Having longer insert lengths was not considered to be vital for contig building or subsequent SNP discovery although it could have compromised genome assembly efforts.

**Figure 2.**
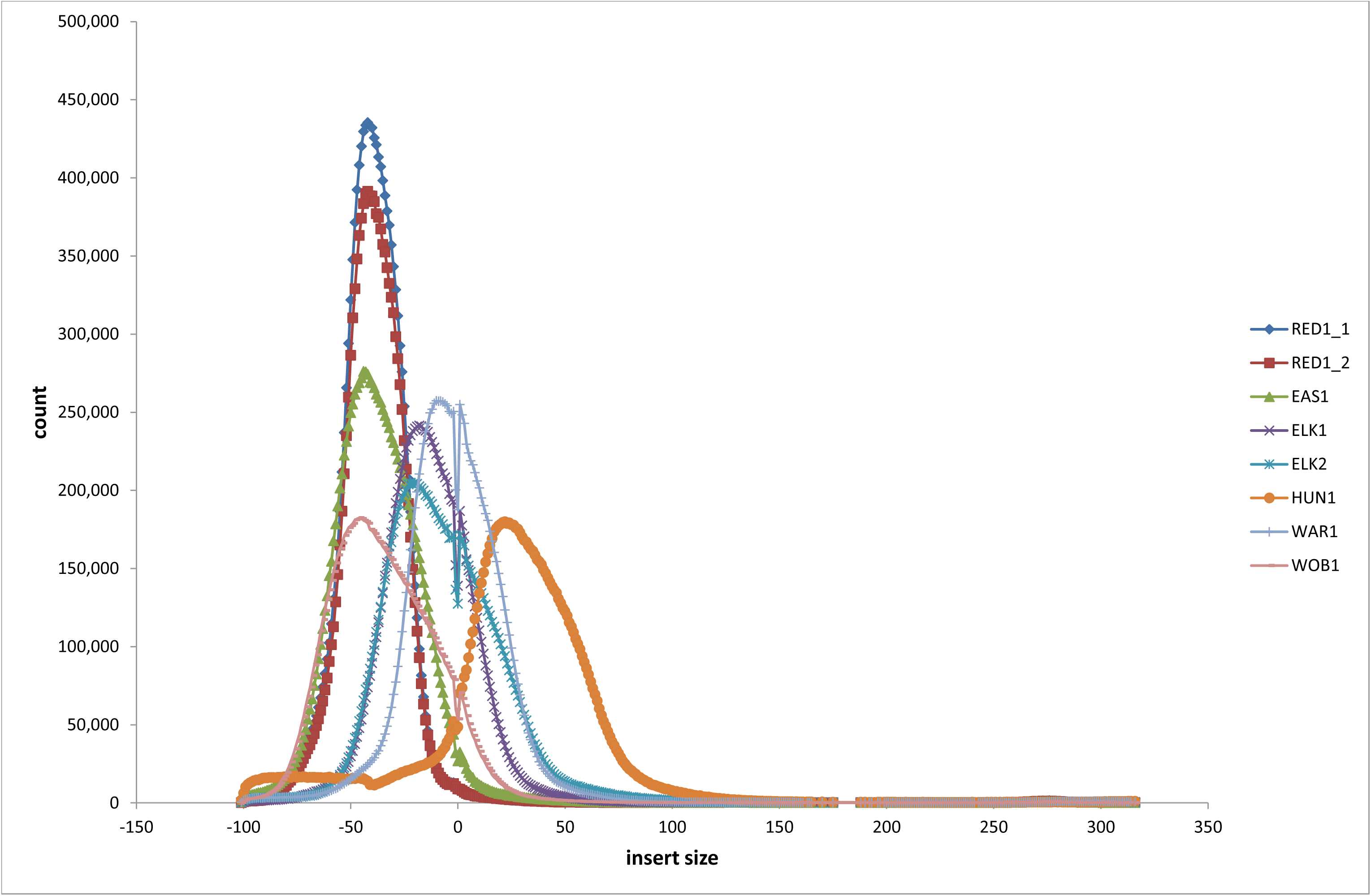
Insert length gap of the reads for the 8 sequence libraries. The count of the insert lengths (distance between the paired sequence reads) is displayed for the eight sequence libraries, Red1 (two libraries), EAS1, ELK1, ELK2, HUN1, WAR1 and WOB1. The minimum value of -100 occurred when the two reads completely overlapped one another. The zero value represents the read counts with no insert and no overlap of the reads. A value of zero means that the fragment size equals exactly two times the read length. Only the first Hungarian’s sequence library provided read pairs that typically were separated by non-sequenced insert DNA. Only rarely were any of the read pairs separated by more than 100 bp insert.

### Assembly and Scaffold-building

The initial Velvet assembly, with a kmer of 31, produced three million contigs with an N50 of 813 bp. In total we assembled 1.6 Gbp, equating to 59% of the cattle genome. Only 59.5% of the bovine reference genome is unique sequence, the rest is Ns and repeats (Adelson et al. 2009).

Cattle and deer share a common Pecoran ancestry and are likely to have diverged 15-20 million years ago (Scott and Janis 1993; Slate et al. 2002; Hassanin and Douzery 2003) this information plus previous mappings of deer sequence to the bovine genome reference (not shown) had suggested that the bovine reference genome would be suitable for comparative genome assembly. Further, whilst this would result in a bovine, rather than deer-ordered genome, only seven large-scale chromosomal red deer/cattle rearrangements had been reported based on karyotype and genetic marker analysis (Slate et al. 2002). The high level of specificity and comprehensive genome coverage on ~99% of the non-repetitive bovine genome confirmed that UMD3.0 had indeed been a suitable reference source.

Changing the order, orientation and spacing during the within-bin scaffolding procedure with the MELD program reduced the number of contigs by 34% and the overall length covered by 8%; it also increased the N50 by 26% to 1021 bp. In general, the assembly procedures were deemed to have worked appropriately because just 13% of NCBI cattle mRNA refseqs (Release 38) did not map to the deer scaffolds; and this was primarily because they were in masked regions

### SNP detection and filtering

Figure 3 shows the distribution of read depths; this closely fits a Poisson distribution, as expected for non-repetitive loci, although it was noted that right hand tail may be slightly higher indicating higher depth reads. The first filtering of SNPs, the depth filter, was applied using this data; SNPs covered by < 3 or >17 reads were removed from further investigation; this resulted in a total of 5.8 million. The numbers of putative SNPs remaining after application of the four more filters, *proximity, class, nucleotide exchange,* and *InfiniumII design threshold* are shown in Table 2. First, application of the proximity filter resulted in the removal of 38% of the SNPs. Next, we observed that 2.4 million of the 4.1 million SNP variants were classified as class A SNPs i.e. both alleles were seen more than once in the entire seven-animal dataset. Another 1.6 million SNPs were class B SNPs, whilst just 31,411 class C SNPs remained. Class A SNPs were considered to be more likely to be reliable for developing SNP markers as non-systematic sequence error is less likely to occur at the same position multiple times. In total across the genome, 3.6 million, or 88% of the SNPs are A/C and A/G exchanges. The distribution of the 4 SNP alternative nucleotide exchanges is shown in Figure 4 for chromosome 1 which was representative of the distribution across entire genome. The A/T and G/C SNPs cannot be converted into *Infinium II* SNP chip markers and were therefore removed from the class A SNP set leaving 2.15 million SNPs. The counts of distances between adjacent class A SNPs, including distances between the outermost SNPs and the ends of their retrospective chromosomes were recorded for this dataset (Figure 5). In summary, 99% of the markers were separated by 10 kb or less, 70% were separated by 1kb or less and 25% of the SNP pairs were located within 100 bp of one another. These values may not be an exact representation of the true distribution of SNP variants in the non-repetitive genome because the proximity filter above is likely to have removed genuine SNPs that were very close to one another. However, inter-SNP distances can be useful for future SNP chip building. These 2.15 million SNPs, and their flanking DNA were submitted to Illumina to put through their *Infinium II* design pipeline; 1.98 million SNPs (>90%) passed the design threshold of 0.8. These SNPs were deemed to be amongst the most suitable SNPs for designing *InfiniumII* SNP chips from.

**Figure 3.**
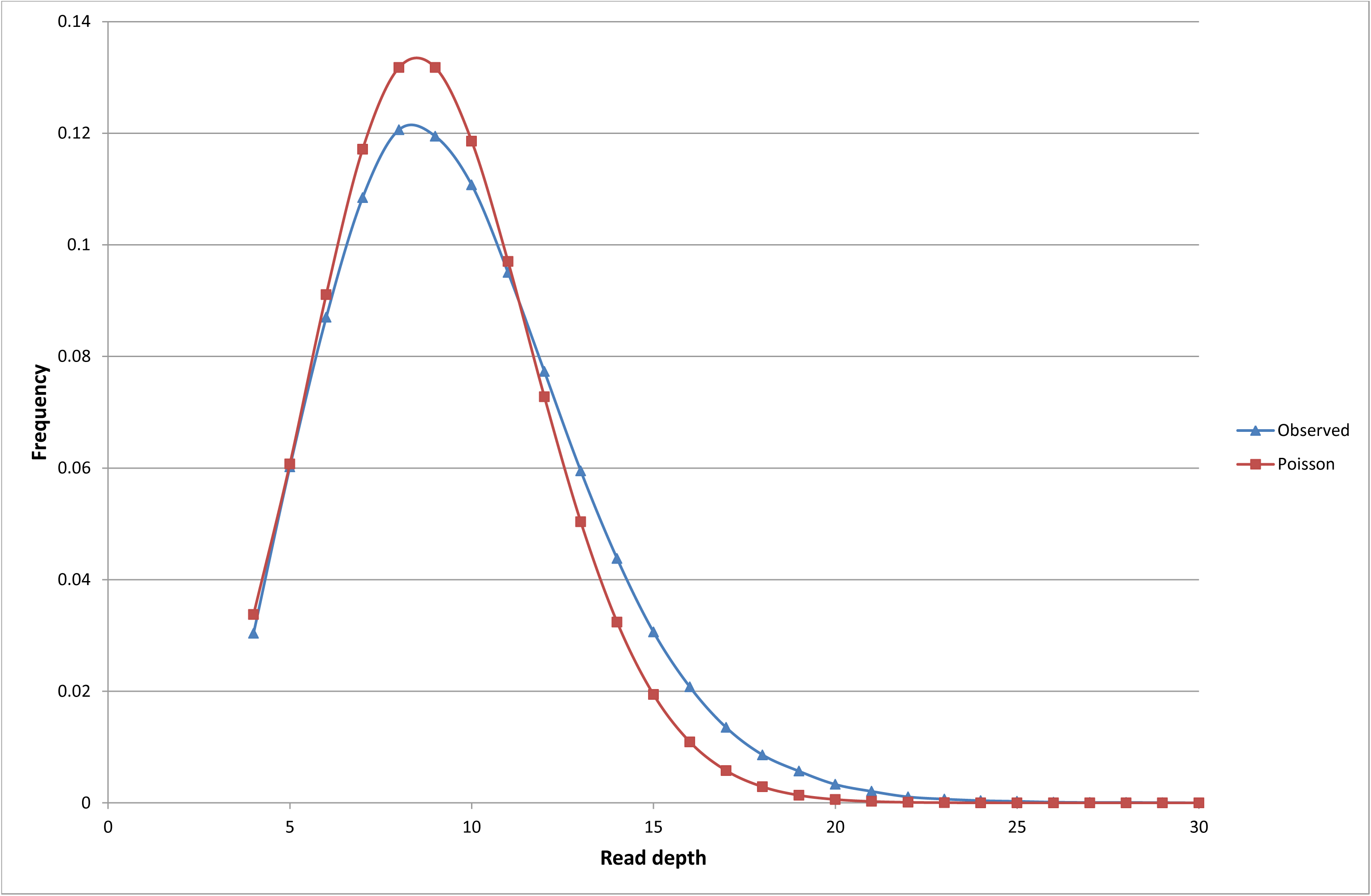
Distribution of Read Depth. The combined frequency of the read depths of the assembled non-repetitive reads of all libraries. The theoretical Poisson distribution for an average depth of nine reads per locus is shown in light grey. The actual, observed distribution is also shown (dark grey).

**Figure 4.**
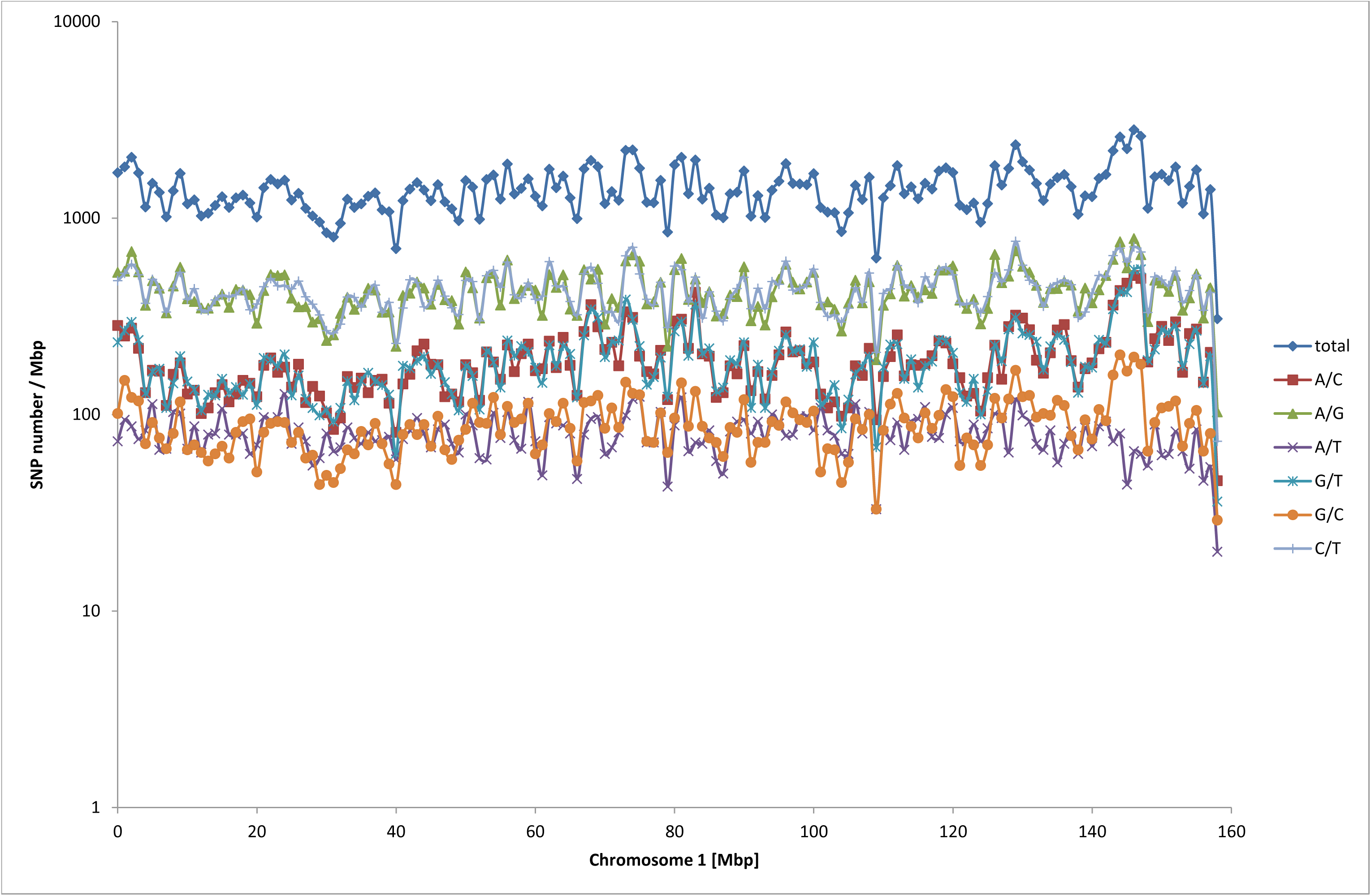
Distribution of alternative SNP types. The SNP count for each consecutive one megabase bin is shown for all putative SNPs that map to bovine chromosome 1. The total count, as well as the count for each motif pair type of i.e. A/T, A/C, A/G and G/C is given.

**Figure 5.**
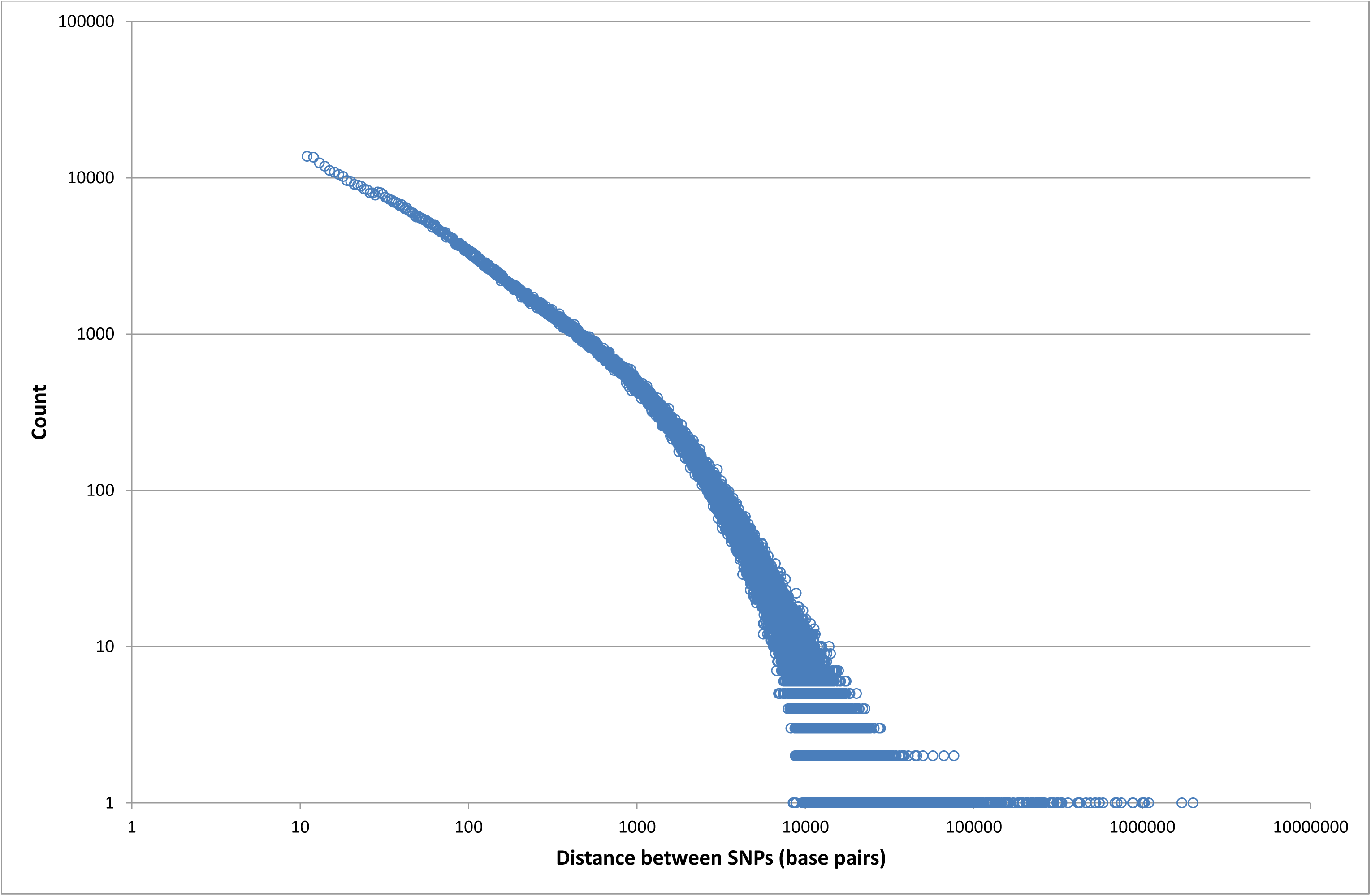
Distances in base pairs between pairs of adjacent SNPs. Counts of the distances in base pairs are shown for pairs of SNPs according to their map positions in UMD3.0. The distances between the start of each chromosome and the first ordered SNP, and between the last ordered SNP and the end of its chromosome are included.

**Table 2.** SNP counts by class and *Infinium* SNP chip type. The numbers of putative SNPs are shown for various SNP classes (A, B or C). To be considered as class A SNPs, both alleles must be observed in more than one individual. Class B SNPs require that only one of the alleles is observed in more than one individual. Class C SNPs occur when both alleles are seen in only one individual. The number of SNPs that would be suitable for *Infiniumll* SNPs chips i.e. that include A/C or A/G motif changes only is given (row 3). The numbers shown in row 4 indicate additional SNPs that would have been suitable for *InfiniumI* technology i.e. that also include A/T and G/C motif changes. The sum of these two rows is given in the Grand Total.

| **SNPs detected** |  |  |  |  |
| --- | --- | --- | --- | --- |
| **SNP Class** | **A** | **B** | **C** | **Grand Total** |
| ***InfiniumII*** | 2,146,571 | 1,451,010 | 27,673 | 3,625,254 |
| ***InfiniumI*** | 289,946 | 195,993 | 3,738 | 489,677 |
| **Grand Total** | 2,436,517 | 1,647,003 | 31,411 | 4,114,931 |

### Presence of SNPs in red deer vs Canadian elk

Genetic distances between *C. e. hippelaphus* “Eastern European”, *C. e. atlanticus* “Western European” and *C. e. canadensis* “Canadian Elk” have been reported using mitochondrial information (Pitra et al. 2004). These data, supported by our own (unpublished) data, indicates that the five United Kingdom and European red deer are more closely related to one another than *B. taurus* is to *B. indicus* i.e. less than a 200,000 year old taxonomic split (Loftus et al. 1994). Using this analogy we might predict that most of the SNPs that are polymorphic in one red deer sub-species will also be polymorphic in the other. However, the Canadian elk is ~3.5 million years distant to the red deer (Pitra et al. 2004) and we would expect a much lower proportion of SNPs to be polymorphic in both elk and red deer. Table 3 shows how many of the two million SNPs were present in (i) Canadian elk only (ii) red deer only, (iii) both elk and red deer and (iv) fixed in one and non-existent in the other. Note that this SNP information should not be used for estimates of across-sub-species polymorphism, because the information from two elk hinds is under-represented compared to the five red animals. Rather, the table highlights the fact that the majority of these SNPs will be of value at least in red deer. It also shows that the number of SNPs that contain one fixed allele exclusively in the reds and the other, exclusively in the elks is high (0.30). Again, caution must be exercised due to low read depth, but this suggests that sub-species-defining SNP alleles may be present.

**Table 3.**
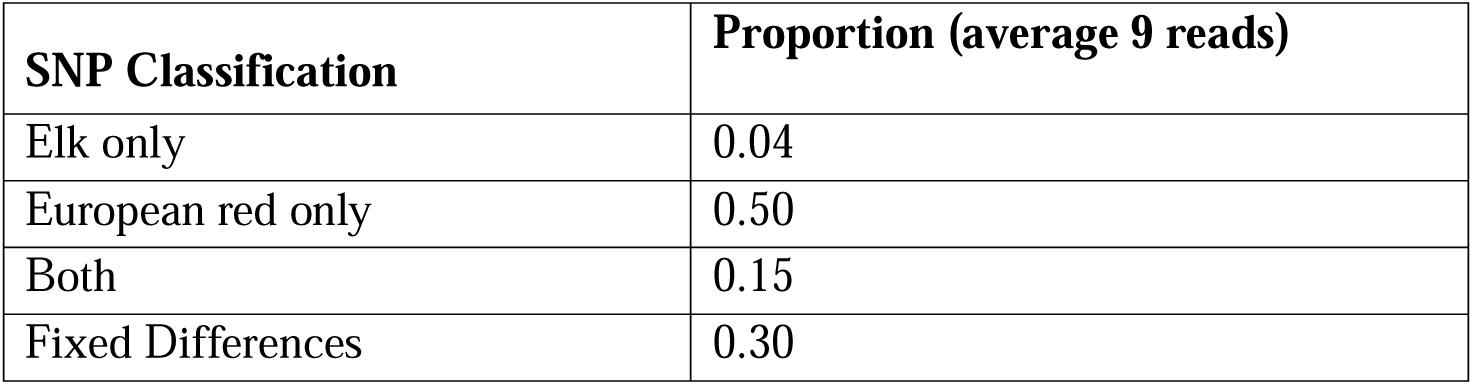
Sequence-derived SNP frequency comparisons across sub-species. The proportions of class A SNPs that were polymorphic amongst the elk animals only, the red deer only and in both red and elk are shown. Putative sub-species distinguishing SNPs that are shown in row 5; here, the allele that is seen and fixed in elk is different to the allele that is fixed in red deer. These values were recorded for SNPs that had an average coverage depth of 9 reads with a minimum depth as low as 3 reads.

| **SNP Classification** | **Proportion (average 9 reads)** |
| --- | --- |
| Elk only | 0.04 |
| European red only | 0.50 |
| Both | 0.15 |
| Fixed Differences | 0.30 |

### Genotypic validation of SNP calls

A set of eight Sequenom multiplexes consisting of 270 SNP markers were genotyped to validate the sequence-based SNP predictions. The average call rate of technically acceptable genotypes for these 270 class A markers was 88%. Thirty three SNPs failed to produce genotypes with acceptable quality criteria; i.e. where “conservative” or “moderate” Sequenom calls occurred in less than 90% of the tested samples. An attrition rate is expected for Sequenom panels containing more than 30 markers in the absence of optimization of the test chemistry. Therefore a call rate of 88% is likely to be a conservatively low estimate of the number of sequence-predicted SNPs that could produce valid markers.

Hardy-Weinberg Equilibrium (HWE) values were calculated and 4.7% of the “acceptable” 237 SNPs did not make the P < 0.05 threshold. As this is close to the 5% level that would be expected by chance, we concluded that there was no evidence for a consistent departure from HWE amongst these markers.

Allelic dropout (ADO), where the parent was homozygous and the confirmed offspring was homozygous for the alternate allele, was observed five times. The number of genotype pairs that ADO could be observed in for this dataset (where one of the pair was genuinely heterozygous and the other was homozygous) was 420. Therefore this type of error rate was estimated to be ~1%.

There was no indication from the genotype profiles that the SNPs were in repetitive regions. Further, there were no occasions when a marker indicated that the individuals were all heterozygous for its two alleles, suggesting that our masking, mapping and filtering processes were effective at minimizing the selection of SNPs in repetitive regions.

<u>Monomorphism and Minor Allele Frequencies</u>: The minor allele frequencies of the 237 validated SNPs are shown in Figure 6 for each breed sample set. It is unsurprising that almost half of the elk SNPs had allele frequencies of less than 5% because no consideration was given to the selection of SNPs that were polymorphic in both elk and red samples. In fact, just 33 of the 237 genotyped SNPs (14%) were polymorphic in the two sequenced elk samples; 27 of these SNPs were also polymorphic in the five sequenced red deer. Comparatively, 168 of the 237 SNPs (71%) were polymorphic amongst the five sequenced red deer. The remaining 36 SNPs were putatively breed-defining SNPs; one allele was observed only in the two elk whilst the alternative allele was observed only in the five red deer. However, none of these 36 SNPs were subsequently shown to be completely fixed in each of the breed reference samples. The relative dearth of elk polymorphic SNPs is assumed to be due to under-representation at the sequence level relative to the red samples rather than a reflection of biological differences between the sub-species. Upon investigation of SNP variation in the two Eastern European red deer, who had a similar combined read depth to the elk samples, it was found that the exact same number of SNPs (33) were polymorphic.

**Figure 6.**
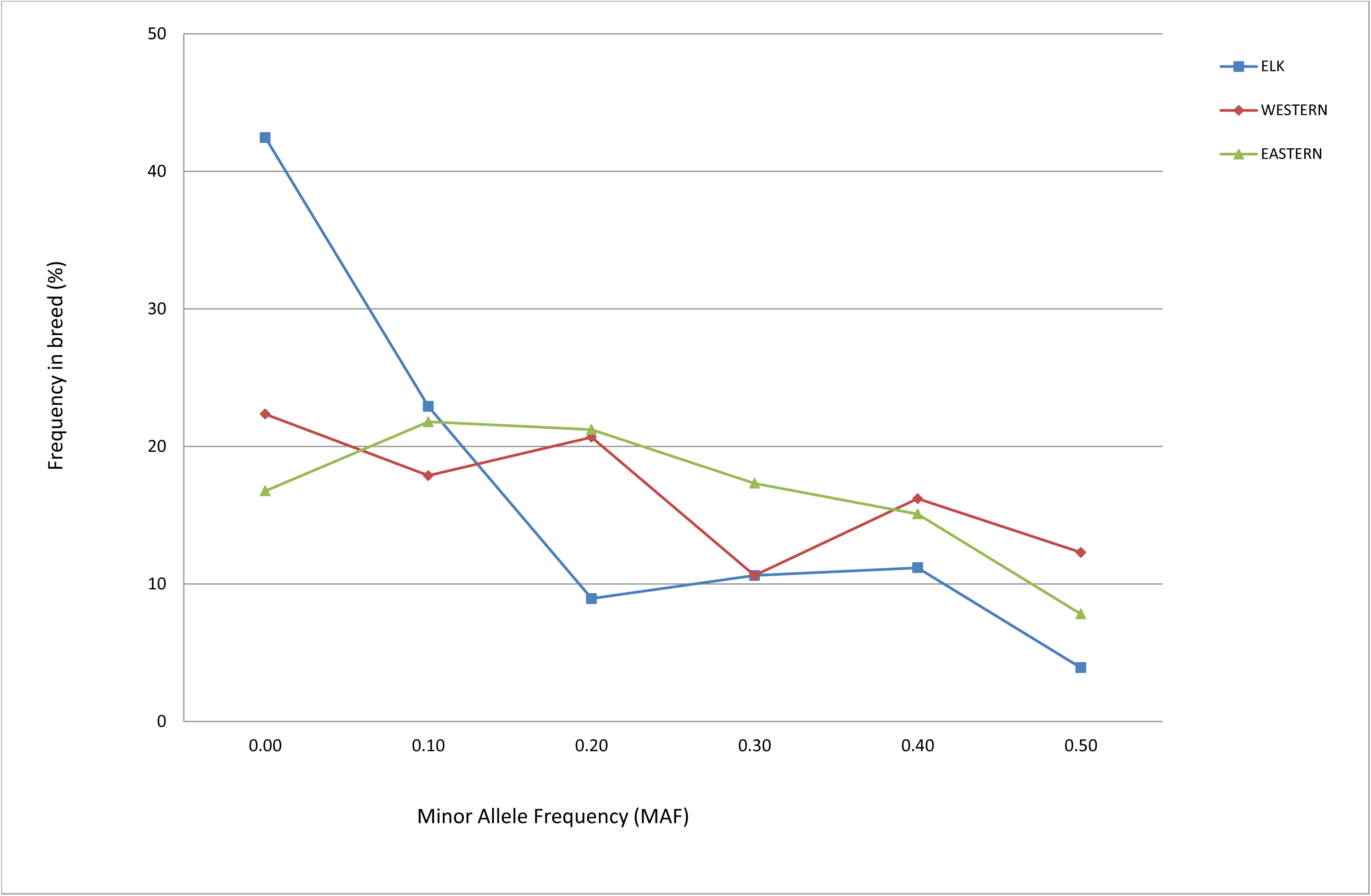
SNP Minor Allele Frequencies in breed reference sample sets. Minor allele frequencies (MAFs) are shown for Western European red (red), Eastern European red (green) and Canadian elk (blue) breed reference sample sets.

Notably, 23 of the putative autosomal SNPs (8.9%) were homozygous in all three breed reference panels and for the original seven sequenced deer that were also included in the genotyping. The seven deer had a total depth, averaging 8.6 per SNP (range 5-13) and 22 of the SNP calls implicated that at least one of the animals was heterozygous. The implication of these discrepancies is that the sequence, bioinformatic approach and/or the genotyping platform, rather than inappropriate population sub-sampling, might have yielded the false SNP calls.

Essentially, the genotyping program implicated that the sequencing approach had been successful for SNP discovery. However, the fact that ~9% of these SNP calls may be false indicates that further improvements in SNP selection may be possible. Several approaches use reduced representational libraries (RRLs) to obtain large numbers of SNPs in mammals (Van Tassell et al. 2008; Kerstens et al. 2009) including, white tailed deer (Seabury et al. 2011). Variations of these approaches, combined with efficient tagging of multiple individuals prior to sequencing add value because they allow for genotyping directly from sequencing (Elshire et al. 2011). However, this study indicates that large numbers of SNPs can be discovered without the need to concentrate on only part of the genome via RRLs. Rather, the adopted approach was shown to be representative of the whole genome. And the data show that millions of SNPs can be identified from 28.4 Gbp of sequence output. Recent technological advances have led to the widespread implementation of the HiSeq2000 and 2500 platforms (Illumina Inc.), which can produce similar outputs to this from just a single lane. The increased cost-effectiveness of sequence generation and the continued production of reference genome assemblies make our approach to SNP discovery more feasible than ever before.

### Mapping deer linkage groups to the UMD3.0 bovine reference sequence

A Pere-David x red deer linkage map was previously aligned to bovine genetic positions but had not been aligned to a bovine genome reference sequence. The mapping of linkage groups to bovine reference genome sequence positions was carried out to confirm the published deer-cattle chromosomal homologies (Slate et al. 2002) and also to confirm that all of the mapping procedures reported in this document were in agreement. Using the Mega BLAST conditions described (see Methods), 113 microsatellites and 123 RFLVs were mapped to UMD3.0. The average marker sequence read length was 547 bp. All marker assignments to bovine chromosomes were in agreement with the predictions described in the literature (Ihara et al. 2004; Slate et al. 2002). Also, 100 of the microsatellites and all of the RFLVs were successfully mapped to the deer chromosomal scaffolds; these deer chromosomal scaffolds had already been assigned UMD3.0 positions during the contig assembly phase. In every case, the direct marker UMD3.0-alignments were in agreement with the corresponding deer contig UMD3.0 positions. In the majority of cases, the syntenic relationships of the deer linkage groups and bovine chromosomes matched those described in a previous study (Slate et al. 2002) with the following exceptions: (i) two of the microsatellite markers (OCAM and CSSM14) were mapped to different bovine chromosomes than had been reported in the deer linkage paper (Slate et al. 2002); at the time of publication, the paper did not have access to the comprehensive bovine linkage map that is now available (Ihara et al. 2004). Subsequent bovine linkage mapping to this higher resolution mapping resource had reassigned the two markers to new bovine linkage map positions; our sequence alignments are in accordance with these most recent linkage map assignments (Ihara et al. 2004). (ii) Nine RFLV sequences mapped ambiguously to multiple chromosomes and their positions could not be unambiguously resolved. Also 14 RFLVs (*LHB, CLTAlike3, FGF2, AT3, HBQ1, TCRBlikel, MYF5like, PRKCB1, NDUFV2like, LHCGR TCRBlike2, CLTAlike2, ASLlike and ANXllikel)* each mapped to a single bovine position but the chromosome was different to the expectations of (Slate et al. 2002). Reciprocal BLAST searches were carried out and in all cases the results were consistent with the original BLAST predictions; that is, the sequence alignments remained ambiguous or were inconsistent with expected locations. The mapping positions of the 14 RFLVs remain ambiguous and unresolved.

The comparative maps, relating the deer genetic map positions (cM) to bovine UMD3.0 positions (Mbp) are shown in Additional file 2. The deer linkage group positions, as defined by (Slate et al. 2002), are shown for the “LG” chromosomes (black). The CELA chromosomal scaffold positions correspond approximately to bovine locations (Mbp, UMD3.0). For every chromosomal match there were at least two links, the minimum required to provide orientation. In general, the maps aligned well, showing consistent marker orders according to expectations from the previous study. There were four occasions where orders of marker pairs, located on deer linkage groups LG2, LG14, LG23 and LG24, differed marginally from expectations. However, all of the marker pairs mapped to the correct region and were either at the ends of their linkage groups and/or located within one Mbp of one another suggesting that these ordering discrepancies may be caused by insufficient linkage mapping resolution.

Deer linkage group names were assigned to the UMD3.0-mapped deer chromosomal scaffold in an order consistent with the bovine genome, but they were additionally assigned a deer linkage group suffix. Whilst this assignment of deer linkage groups to the deer chromosomal scaffold is useful, it is recognized that renaming may be required as new deer assemblies and/or mapping programs are advanced.

Bovine chromosomes 1, 2, 5, 6, 8 and 9 are exceptional because they are homologous to the deer *fission* chromosomes; not all contigs that map to them can be placed unambiguously to deer linkage groups. Deer linkage groups and corresponding bovine sequence locations are shown for the 28 microsatellites and 37 RFLVs that mapped to these six bovine and twelve deer chromosomes (Table 4). Every marker mapped to the expected chromosomal position according to the published linkage map, with the exception of INHBB. This marker mapped to the expected cattle chromosome (UMD3_3), but its physical location placed it in the middle of the deer linkage group eight markers, not at the end of linkage group 33, as predicted in the literature (Slate et al. 2002). Given that every linkage group was represented by at least two markers, there was sufficient information to assign linkage group nomenclature to contigs, except where they mapped between the last microsatellite of CELA_A and the first microsatellite from CELA_B. For example, chromosomal scaffold segments that mapped between positions 0 Mbp and 19 Mbp on bovine chromosome 1 were assigned to deer linkage group 31. Contigs that mapped between 60 Mbp and the terminal end of the same bovine chromosome were assigned to deer linkage group 19. But chromosomal scaffold segments that mapped between 19 Mbp and 60 Mbp on bovine chromosome 1 could not be assigned unambiguously to either linkage group although it is recognized that they are likely to map to either deer linkage group 31 or 19. Therefore in this case, the unresolved gap which most likely harbors a breakpoint was 41 Mbp. On average, based on these data alone, the chromosomal breakpoint positions were located within unresolved regions averaging 29.5 Mbp. In order to locate the breakpoints more precisely, a denser or more targeted genetic mapping program, or a comprehensive deer assembly aimed to significantly improve the scaffolding building would be required.

**Table 4.**
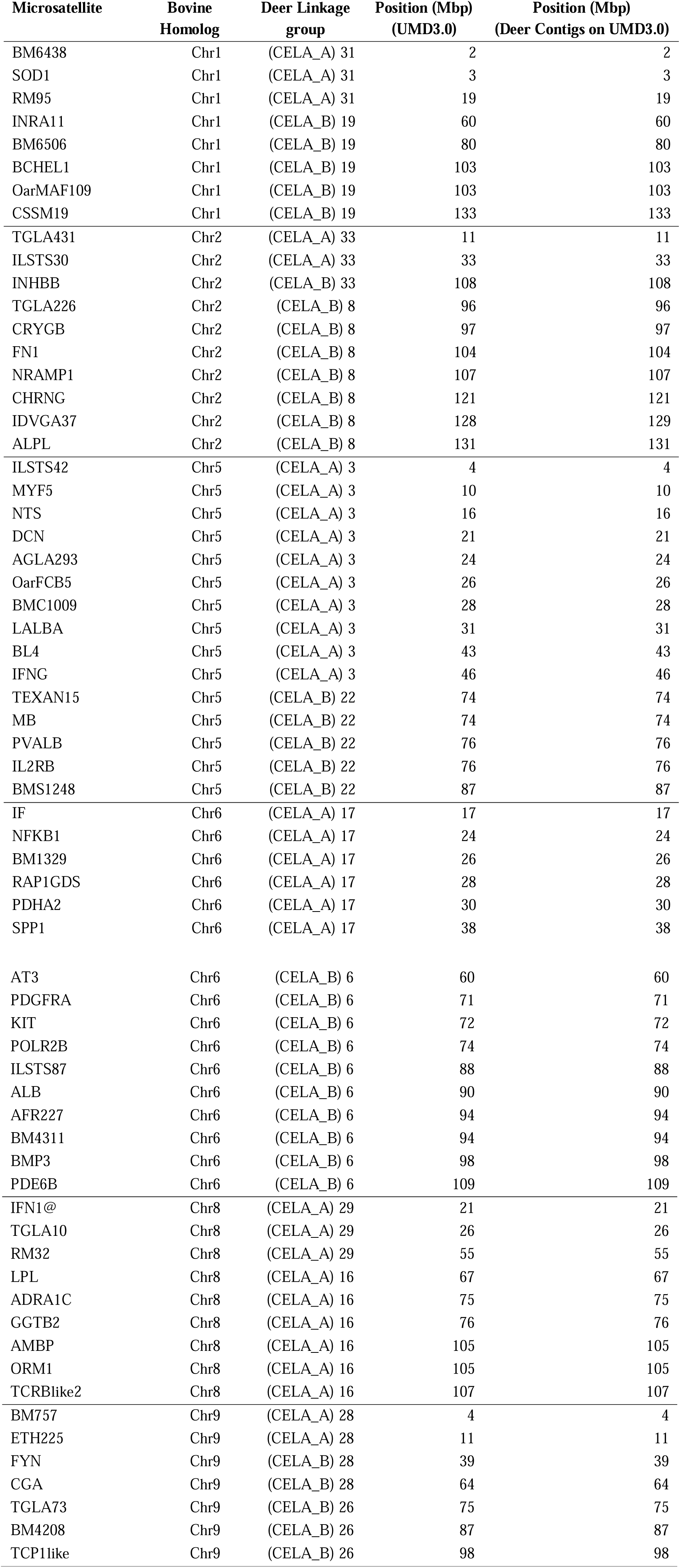
Mapping locations of markers mapped to deer fission chromosomes. Twenty eight microsatellites and 37 RFLVs were mapped via BLAST alignments to the UMD3.0 cattle reference genome positions for six bovine chromosomes. These chromosomes corresponded to 12 deer linkage groups representing chromosomes that had undergone fission. The markers were also mapped to deer contigs; column 5 shows these contigs’ UMD3.0 positions. The markers are shown listed in genetic order, except for CSSM19 which maps distal to BM6506 on deer linkage group 19 and is situated inside a reported inversion (Slate et al. 2002)

### Conclusion

The described procedure revealed that, from just 28 Gbp of sequence output (i.e. less than the amount of sequence generated by a single HiSeq2000 lane today) 1.8 million cervid *InfiniumII*-compatible SNPs could be identified. The protocol initially utilized repeat masking techniques and resources to reduce the complexity of the assembly. Homology to a bovine reference genome reduced complexity during contig assembly phase even further, providing manageable 1 Mbp blocks of data. Not only did the approach provide very high coverage of non-repetitive DNA, but subsequent genotyping results showed that the SNPs had not been inadvertently selected from repetitive regions. Further, the genotyping confirmed that the entire procedure, including the application of four SNP-selection filters had been sufficient to produce valid and useful SNPs.

The described protocol was designed to identify SNPs rather than to build a deer-ordered reference genome and the chromosomal scaffolds were based on bovine order. As ~99% of the non-repetitive bovine genome is covered by this deer sequence, the resource would be valuable for including in a future cervid reference genome assembly program.

## AUTHORS’ CONTRIBUTIONS

PF secured funding, was responsible for the project, contributed to trial design, analyzed the genotyping results, contributed to the mapping and wrote the manuscript. RB carried out the bioinformatics, including assembly, SNP detection and comparative mapping, contributed to project design and assisted in the writing of the manuscript. AM and RS provided IT support and bioinformatic analysis. JW sourced the animals and provided their pedigrees. MB carried out all of the lab work excluding the sequencing. CL oversaw the collaborative sequencing component and advised on sequence trial design. SJR oversaw manuscript preparation and submission and contributed to the analysis. JM contributed to trial design, carried out bioinformatic analysis and summary statistics and contributed to the manuscript preparation.

## ACKNOWLEDGEMENTS

The authors would like to acknowledge financial support from Landcorp Farming Ltd, with particular thanks to Dr Geoff Nicoll, and Illumina Inc. that enabled the sequencing program to be carried out. Additional financial support was provided by DEEResearch, which is a joint venture between AgResearch and levy-funded Deer Industry New Zealand (DINZ), for the sequence analysis component.

## Additional files

### Additional file 1

Pedigrees of sequenced animals

### Additional file 2

Comparative Mapping of Deer Linkage groups, LG (cM) & bovine chromosome positions, CELA (Mbp)

### Additional file 3

Draft deer genome. Softmasked Fasta files. Available at http://sheephapmap.org/BRAUNINGDEER/deer_chromosomes_on_UMD3.tar.gz

## REFERENCES

Adelson, D.L., J.M. Raison, and R.C. Edgar, 2009 Characterization and distribution of retrotransposons and simple sequence repeats in the bovine genome. Proc Natl Acad Sci U S A 106 (31):12855–12860.

Dalrymple, B.P., E.F. Kirkness, M. Nefedov, S. McWilliam, A. Ratnakumar et al., 2007 Using comparative genomics to reorder the human genome sequence into a virtual sheep genome. Genome Biology 8 (7).

De Roos, A.P.W., C. Schrooten, R.F. Veerkamp, and J.A.M. van Arendonk, 2011 Effects of genomic selection on genetic improvement, inbreeding, and merit of young versus proven bulls. Journal of Dairy Science 94 (3):1559–1567.

Deer Industry New Zealand, http://deernz.org.nz/

Elshire, R.J., J.C. Glaubitz, Q. Sun, J.A. Poland, K. Kawamoto et al, 2011 A Robust, Simple Genotyping-by-Sequencing (GBS) Approach for High Diversity Species. PLoS ONE6 (5):e19379.

Fisher, P.J., B. Malthus, M.C. Walker, G. Corbett, and R.J. Spelman, 2009 The number of single nucleotide polymorphisms and on-farm data required for whole-herd parentage testing in dairy cattle herds. Journal of Dairy Science 92 (1):369–374.

Frkonja, A., B. Gredler, U. Schnyder, I. Curik, and J. Sölkner, 2012 Prediction of breed composition in an admixed cattle population. Animal Genetics 43 (6):696–703.

Genetic Information Research Institute, http://www.girinst.org/

Hassanin, A., and E.J.P. Douzery, 2003 Molecular and morphological phylogenies of Ruminantia and the alternative position of the Moschidae. Systematic Biology 52 (2):206–228.

Hayes, B.J., and M.E. Goddard, 2008 Technical note: Prediction of breeding values using marker-derived relationship matrices. Journal of Animal Science 86 (9):2089–2092.

Holstein Canada Home Page, https://www.holstein.ca/

Ihara, N., A. Takasuga, K. Mizoshita, H. Takeda, M. Sugimoto et al., 2004 A Comprehensive Genetic Map of the Cattle Genome Based on 3802 Microsatellites. Genome Research 14 (10a):1987–1998.

Illumina Paired-End Sequencing Information, http://www.illumina.com/technology/next-generation-sequencing/paired-end-sequencingassay.html

Kerstens, H.H.D., R.P.M.A. Crooijmans, A. Veenendaal, B.W. Dibbits, T.F.C. Chin-A-Woeng et al, 2009 Large scale single nucleotide polymorphism discovery in unsequenced genomes using second generation high throughput sequencing technology: Applied to Turkey. BMC Genomics 10: 479.

Kijas, J.W., J.A. Lenstra, B. Hayes, S. Boitard, L.R. Neto et al., 2012 Genome-wide analysis of the world’s sheep breeds reveals high levels of historic mixture and strong recent selection. PLoS Biology 10 (2).

Kuehn, L.A., J.W. Keele, G.L. Bennett, T.G. McDaneld, T.P.L. Smith et al., 2011 Predicting breed composition using breed frequencies of 50,000 markers from the US Meat Animal Research Center 2,000 bull project. Journal of Animal Science 89 (6): 1742–1750.

Loftus, R.T., D.E. MacHugh, D.G. Bradley, P.M. Sharp, and P. Cunningham, 1994 Evidence for two independent domestications of cattle. Proc Natl Acad Sci U S A 91 (7):2757–2761.

Meuwissen, T.H.E., B.J. Hayes, and M.E. Goddard, 2001 Prediction of Total Genetic Value Using Genome-Wide Dense Marker Maps. Genetics 157 (4): 1819–1829.

Montgomery, G.W., and J.A. Sise, 1990 Extraction of DNA from sheep white blood cells. New Zealand Journal of Agricultural Research 33 (3):437–441.

Pitra, C., J. Fickel, E. Meijaard, and C.P. Groves, 2004 Evolution and phylogeny of old world deer. Molecular Phylogenetics and Evolution 33 (3):880–895.

Pryce, J.E., and H.D. Daetwyler, 2012 Designing dairy cattle breeding schemes under genomic selection: A review of international research. Animal Production Science 52 (2–3):107–114.

Richter, D.C., F. Ott, A.F. Auch, R. Schmid, and D.H. Huson, 2008 MetaSim - A sequencing simulator for genomics and metagenomics. PLoS ONE 3 (10).

Scott, K., and C. Janis, 1993 Relationships of the Ruminantia (Artiodactyla) and an analysis of the characters used in ruminant taxonomy, pp. 282–302 in Mammal Phylogeny: Placentals, edited by F.S. Szalay, M.J. Novacek and M.C. McKenna. Springer Verlag.

Seabury, C.M., E.K. Bhattarai, J.F. Taylor, G.G. Viswanathan, S.M. Cooper et al., 2011 Genome-Wide Polymorphism and Comparative Analyses in the White-Tailed Deer (*Odocoileus virginianus*): A Model for Conservation Genomics. PLoS ONE 6 (1):e15811.

Slate, J., T.C. Van Stijn, R.M. Anderson, K. Mary McEwan, N.J. Maqbool et al., 2002 A deer (subfamily cervinae) genetic linkage map and the evolution of ruminant genomes. Genetics 160 (4):1587–1597.

Smit, A.F.A., R. Hubley, and P. Green, 1996–2010 RepeatMasker Open-3.0.

Van Tassell, C.P., T.P.L. Smith, L.K. Matukumalli, J.F. Taylor, R.D. Schnabel et al., 2008 SNP discovery and allele frequency estimation by deep sequencing of reduced representation libraries. Nature Methods 5 (3):247–252.

Zerbino, D.R., and E. Birney, 2008 Velvet: Algorithms for de novo short read assembly using de Bruijn graphs. Genome Research 18 (5):821–829.

Zhang, Z., S. Schwartz, L. Wagner, and W. Miller, 2000 A greedy algorithm for aligning DNA sequences. Journal of Computational Biology 7 (1–2):203–214.

Zimin, A.V., A.L. Delcher, L. Florea, D.R. Kelley, M.C. Schatz et al., 2009 A whole-genome assembly of the domestic cow, Bos taurus. Genome Biology 10 (4).

